# Modulation of functional networks related to the serotonin neurotransmitter system by citalopram: evidence from a multimodal neuroimaging study

**DOI:** 10.1101/2022.10.20.512503

**Authors:** Daphne E Boucherie, Liesbeth Reneman, Jan Booij, Daniel Martins, Ottavia Dipasquale, Anouk Schrantee

## Abstract

**Background:** Selective serotonin reuptake inhibitors (SSRIs) potentiate serotonergic neurotransmission by blocking the serotonin transporter (5-HTT), but the functional brain response to SSRIs involves neural circuits beyond regions with high 5-HTT expression. Currently, it is unclear whether and how changes in 5-HTT availability after SSRI administration modulate brain function of key serotoninergic circuits, including those characterized by high availability of serotonin 1A receptors (5-HT1AR).

**Aim:** We investigated the association between 5-HTT availability and 5-HTT- and 5-HT1AR-enriched functional connectivity (FC) after an acute citalopram challenge.

**Methods:** We analyzed multimodal data from a dose-response, placebo-controlled, double-blind study, in which 45 healthy women were randomized into three groups receiving placebo, a low (4 mg), or high (16 mg) oral dose of citalopram. Receptor-Enhanced Analysis of functional Connectivity by Targets was used to estimate 5-HTT- and 5-HT1AR-enriched FC from resting-state and task-based fMRI. 5-HTT availability was determined using [^123^I]FP-CIT single-photon emission computerized tomography.

**Results:** 5-HTT availability was negatively correlated with resting-state 5-HTT-enriched FC, and with task-dependent 5-HT1AR-enriched FC. Our exploratory analyses revealed lower 5-HT1AR-enriched FC in the low dose group compared to the high dose group at rest and the placebo group during the task.

**Conclusions:** Taken together, our findings provide evidence for differential links between 5-HTT availability and brain function within 5-HTT and 5-HT1AR pathways and in context- and dose-dependent manner. As such, they support a potential pivotal role of the 5-HT1AR in the effects of citalopram on the brain and add to its potential as a therapeutic avenue for mood and anxiety disturbances.

## Introduction

Selective serotonin reuptake inhibitors (SSRIs), such as citalopram, are frequently used to treat depression and anxiety disorders (Baldwin et al., 2016; Hieronymus, Emilsson, Nilsson, & Eriksson, 2016). Despite their frequent prescription, approximately one third of patients with depression (Fava & Davidson, 1996) and anxiety (Reinhold & Rickels, 2015) fails to respond to this treatment. Uncovering mechanisms that could account for interindividual differences in treatment response is thus paramount to inform the development of new and more effective therapeutic approaches. However, the current lack of a full mechanistic understanding of the effects of SSRIs on the human brain has marred substantial advances in this direction.

While the exact mechanisms underlying the therapeutic effects of SSRIs remain contentious, the primary pharmacological mode of action entails blocking the serotonin (5-HT) transporter (5-HTT) (Meyer et al., 2004), resulting in increased extracellular levels of 5-HT. Acute SSRI administration thereby results in activation of both the postsynaptic serotonin-1A receptor (5-HT1AR) in projection sites and 5-HT1A autoreceptors of the dorsal raphe nucleus (DRN) (Barnes & Sharp, 1999). Activation of the latter results in hyperpolarization, thereby diminishing 5-HT release from the 5-HT nerve terminals in the synapses and inhibiting neuronal activity (Artigas, 2013). Consecutively, chronic SSRI treatment is thought to induce desensitization of DRN 5-HT1A autoreceptors, thereby decreasing their inhibitory effect and allowing increased 5-HT binding to postsynaptic 5-HT receptors (El Mansari, Sánchez, Chouvet, Renaud, & Haddjeri, 2005; Hervás et al., 2001; Riad, Watkins, Doucet, Hamon, & Descarries, 2001). As such, 5-HT1ARs are thought to play a pivotal role in controlling 5-HT neuromodulation in the brain and have been suggested to contribute to the antidepressant properties of SSRIs (Artigas, 2013; Carhart-Harris & Nutt, 2017).

Our current knowledge of the effects of SSRIs on the human brain has greatly benefited from the use of non-invasive neuroimaging techniques. Particularly, the combination of functional magnetic resonance imaging (fMRI) with an acute pharmacological challenge, i.e. pharmacological MRI (phMRI), has offered an unprecedented opportunity to investigate acute drug effects on brain function at the systems level, which is essential for understanding the psychopharmacological effects of drugs affecting widespread neuromodulatory systems such as 5-HT. Studies employing phMRI at rest have demonstrated that SSRIs modulate activity and connectivity both in brain regions with high 5-HTT density and in their projection areas, likely reflecting downstream effects via 5-HT (and other neurotransmitter) receptors. For example, SSRI administration has been found to alter connectivity within and with the Default Mode Network (DMN) (Arnone et al., 2018; Klaassens et al., 2015; McCabe et al., 2011; Schrantee, Lucassen, Booij, & Reneman, 2018; van de Ven, Wingen, Kuypers, Ramaekers, & Formisano, 2013), which has frequently been associated with rumination in depression (Berman et al., 2011). Furthermore, task-based studies using emotion recognition paradigms have frequently reported altered activation in (networks encompassing) the limbic system in response to SSRIs (Arce, Simmons, Lovero, Stein, & Paulus, 2008; Del-Ben et al., 2005; Grady et al., 2013; Selvaraj et al., 2018; Windischberger et al., 2010), which has been implicated in aberrant emotion regulation in depressed patients (Anand et al., 2005). However, how engagement of 5-HT molecular targets, including the 5-HTT and 5-HT1AR, might contribute to these circuit-level effects remains poorly understood.

Multimodal studies combining phMRI with molecular imaging have shed further light on the involvement of specific 5-HT receptors in modulating FC and SSRI-induced changes in brain function. For instance, one study found that 5-HTT availability following citalopram administration was associated with FC between the DMN and cortical regions (Schrantee et al., 2018), highlighting a prominent role of 5-HTT in modulating the effects of SSRIs on DMN connectivity. Moreover, Hahn et al., (2012) emphasized the importance of 5-HT1AR in regulating DMN connectivity by demonstrating diverse and area-specific patterns of associations between the FC of the posterior cingulate cortex (PCC), a core node of the DMN, and 5-HT1A autoreceptor and local heteroreceptor binding. Noteworthy, 5-HT1AR has also been hypothesized to be a key modulator of neural circuits underlying emotional processing in task-based fMRI studies, as evidenced by associations between DRN 5-HT1AR binding and (changes in) amygdala activation during emotion processing (Fisher, Meltzer, Ziolko, Price, & Hariri, 2006; Selvaraj et al., 2018). Given the prominent role of 5-HT1AR in regulating the 5-HT system, imaging studies dissecting the pharmacodynamics of SSRIs in the brain and its correspondent interindividual variability should incorporate concomitant investigations of the relation between post-drug increases in 5-HT and modulations of neural circuits associated with specific 5-HT receptor subtypes, such as 5-HT1AR. Crucially, both resting-state and task-based paradigms seem to capture interindividual variation in molecular target bioavailability on brain function, and could provide new insights on the context-dependent modulation of 5-HT brain networks by SSRIs.

To this end, we conducted a multimodal study combining single-photon emission computerized tomography (SPECT) with Receptor-Enriched Analysis of functional Connectivity by Targets (REACT) (Dipasquale et al., 2019) to investigate how acute citalopram-induced variations in 5-HTT availability relate to FC of 5-HTT- and 5-HT1AR-enriched functional networks at rest and during an emotional face-matching task. Forty-five healthy women were randomized into three treatment groups receiving placebo, a low (4mg) or high (16mg) oral dose of citalopram; SPECT, resting state fMRI (rs-fMRI), and task-based fMRI (tb-fMRI) were acquired within the same participants to allow between-modality associations across treatment. In line with previous findings, we hypothesized an inverse relationship between 5-HTT availability and 5-HTT- and 5-HT1AR-enriched FC both at rest and during the task. In accordance with previous studies showing that amygdala activation during emotion recognition is associated with DRN 5-HT1AR binding (Fisher et al., 2006; Selvaraj et al., 2018), we expected a stronger association between 5-HTT availability in task-dependent FC and the 5-HT1AR-enriched functional network compared to the 5-HTT-enriched functional network.

## Materials and methods

### Participants

Forty-five healthy female volunteers (HC) were enrolled in the study. Baseline demographics are shown in Table 1. Exclusion criteria included a history of a chronic neurological/psychiatric disorder, family history of sudden heart failure, current use of psychostimulant medication, abnormal electrocardiogram, excessive consumption of alcohol (>21 units/week), caffeine (>8 cups/day), or nicotine (>15 cigarettes/day), and neuroimaging contraindications. The Mini-International Neuropsychiatric Interview Plus was used to screen for psychiatric illnesses and drug abuse. To avoid confounding effects of the hormonal cycle, participants were required to use hormonal contraception. The study protocol was approved by the medical ethics committee of the Academic Medical Center in Amsterdam and in accordance with the Declaration of Helsinki. All subjects gave written informed consent. Prior analyses on the same sample assessed the relation between 5-HTT availability and DMN FC (Schrantee et al., 2018) and dose-dependent effects on 5-HTT availability and arterial spin labeling phMRI (Schrantee et al., 2019). An *a priori* sample size calculation was conducted for the primary outcome measures of the study, which is discussed in more detail in Schrantee et al. (2019).

**Table 1.** Participant demographics and baseline group differences. BAI: Beck Anxiety Inventory; BDI: Beck Depression Inventory. IQ as assessed by the Dutch Adult Reading Test (Schmand, Bakker, Saan, & Louman, 1991). Group differences in baseline characteristics were determined using one-way ANOVAs. Data is represented as mean (SD).

|  | Full sample | Placebo | Low dose | High dose | Group differences |
| --- | --- | --- | --- | --- | --- |
| Sample size | n=45 | n=15 | n=15 | n=14 | ANOVA |
| Citalopram dose (mg) | - | 0 | 4 | 16 | p-value |
| Age (years) | 21.58 (2.4) | 21.3 (2.2) | 22.1 (3.2) | 21.4 (2.0) | 0.66 |
| Weight (kg) | 65.8 (10.5) | 67.1 (10.7) | 62.0 (12.1) | 68.4 (7.7) | 0.22 |
| IQ | 110.8 (7.3) | 113.7 (8.3) | 108.7 (6.6) | 109.9 (6.2) | 0.15 |
| BDI (sum score) | 3.1 (4.1) | 3.5 (4.7) | 3.8 (5.0) | 1.8 (1.5) | 0.37 |
| BAI (sum score) | 4.5 (4.3) | 4.0 (4.6) | 5.0 (3.4) | 4.4 (5.4) | 0.82 |
| Alcohol consumption<br>(glasses/week) | 4.4 (2.9) | 4.0 (2.2) | 4.3 (2.7) | 5.3 (4.1) | 0.65 |

### Study design

We used a placebo-controlled, dose-response, double-blind design (Figure 1). Following a baseline SPECT scan two hours post-injection of [^123^I]N-ω-fluoropropyl-2β-carbomethoxy-3β-(4iodophenyl)nortropane ([^123^I]FP-CIT; data not shown; approximately 110 MBq; specific activity > 750 MBq/nmol; radiochemical purity > 98%, produced according to GMP criteria at GE Healthcare, Eindhoven, the Netherlands), participants were randomly assigned to one of three treatment groups: placebo (n=15), low dose (4mg; n=15), or high (clinical) dose (16mg; n=15) of citalopram (oral solution, 16mg equivalent to 20mg in tablet form, Lundbeck). These doses were selected as they correspond to 5-HTT occupancy levels of 0%, 40%, and 80%, respectively (Klein et al., 2006). Participants underwent a second SPECT scan to assess 5-HTT occupancy by placebo or citalopram three hours after drug intake (i.e. six hours after [^123^I]FP-CIT administration), followed by an MRI scan one hour later, which included a rs-fMRI and a tb-fMRI scan during an emotional face-matching task. It has previously been shown that Cmax uptake of citalopram to SERT is stable from 3 hrs to 22-27 hrs onward (Pirker et al., 1995)

**Figure 1.**
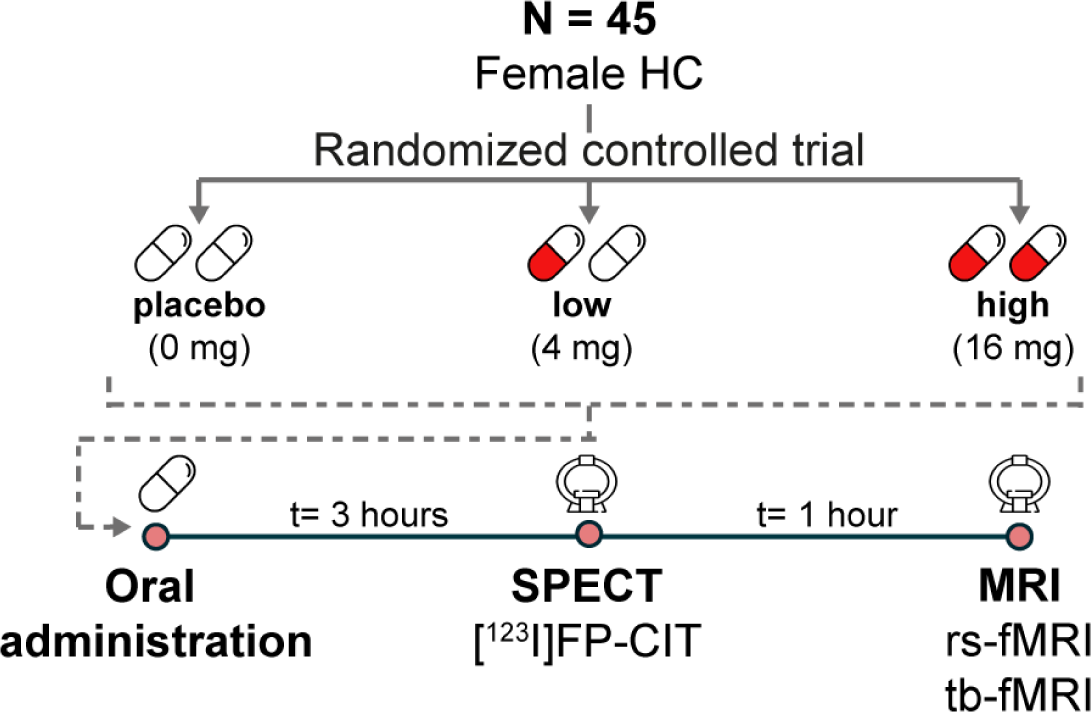
Experimental design. HC: healthy controls; mg: milligram; t: time; SPECT: single-photon emission computerized tomography; [^123^I]FP-CIT: [^123^I]N-ω-fluoropropyl-2β-carbomethoxy-3β-(4-iodophenyl) nortropane; BR: ratio of specific to non-specific binding; fMRI: functional Magnetic Resonance Imaging, rs-fMRI: resting-state fMRI; tb-fMRI: task-based fMRI

### SPECT acquisition and analysis

SPECT scans were acquired using a brain-dedicated InSPiraHD SPECT camera (Neurologica, Boston, USA) with scan parameters: matrix = 121 × 121; slice thickness = 4 mm; acquisition time per slice = 180 s; energy window = 159 keV (with 20% lower and upper boundaries), as described previously (Schrantee et al., 2018, 2019). In short, an iterative reconstruction algorithm was used to reconstruct the data into 3D images, as described earlier (Schrantee et al., 2018, 2019). These 3D SPECT images were co-registered to the individual T1w MRI scan and a region-of-interest (ROI) analysis was performed with ROI masks extracted from the individual T1w image using FreeSurfer. As the radioligand [^123^I]FP-CIT has been shown to bind with high affinity primarily to the dopamine transporter in the striatum, and primarily to the 5-HTT in extrastriatal areas such as the thalamus (Booij et al., 2007), we determined 5-HTT binding in the thalamus with the cerebellum as a reference region reflecting non-specific binding. The ratio of specific to non-specific binding (BR) after drug administration were calculated, reflecting 5-HTT availability (here, a lower BR reflects a higher occupancy and thus a lower availability of the 5-HTT).

### MRI acquisition and preprocessing

MRI data was acquired using a 3.0T Ingenia scanner (Philips, Best, The Netherlands) with a 32-channel receive-only head coil. A high-resolution 3D T1w scan was obtained (TR/TE=3195/7ms; FOV=256×256×180; voxel size=1×1×1mm; flip angle=9°). Rs-fMRI data were acquired using a gradient-echo echo-planar imaging sequence: TR/TE=2150/27ms; FOV=240×240×131mm, voxel size=3×3×3mm; gap=0.3mm; flip angle=76.2°; dynamics=240 (∼9min). Participants were instructed to keep their eyes open, focus on a fixation cross, and let their mind wander. Tb-fMRI data were acquired with the following parameters: TR/TE=2300/30ms, FOV=220×220×117mm; resolution=2.3×2.3×3mm, 39 sequential slices, flip angle=80°, dynamics=70 (∼3min). The pseudo-randomized emotional face-matching fMRI paradigm (adapted from Hariri, Tessitore, Mattay, Fera, & Weinberger, 2002), consisted of a blocked design with emotional stimuli displaying angry and fearful faces (*faces*) and neutral stimuli displaying ellipses assembled from scrambled faces (*shapes*). Two blocks of emotional stimuli were interleaved with three neutral blocks, with each 30-s block containing six 5-s trials. Three stimuli were displayed simultaneously, and individuals had to choose which of the lower two stimuli showed the same emotion or orientation as the target stimulus above, for the emotional and neutral trials respectively.

All fMRI data were preprocessed using fMRIPREP v.20.0.6 (Esteban et al., 2019), which is based on *Nipype* 1.4.2 (Gorgolewski et al., 2011). T1-weighted scans were normalized to MNI space. Motion correction (FLIRT), distortion correction (*fieldmap-less*), and T1-weighted coregistration were performed as part of the functional data preprocessing. To generate non-aggressively denoised data, Independent Component Analysis-based Automatic Removal Of Motion Artifacts was employed. Data were spatially smoothed (6mm FWHM; FSL/FEAT v.6.0.0). See Supplementary Methods for further details. Following preprocessing with fMRIPREP, white matter and cerebrospinal fluid (obtained from fMRIPREP) were regressed out of the main signal using FSL v6.0.0, followed by high pass-filtering (100s). One subject with insufficient scan quality was removed from all analyses and three subjects exhibiting high motion (mean framewise displacement>0.25mm) were excluded from the rs-fMRI analysis.

### Target-enriched FC at rest

A high-resolution in vivo positron emission tomography (PET) atlas of the distribution density of 5-HTT and 5-HT1AR (Beliveau et al., 2017) was used for the REACT analysis (Figure 2). In a two-step multiple linear regression employing REACT (Dipasquale et al., 2019) and FSL, the PET atlas was used as a molecular template to estimate the 5-HTT- and 5-HT1AR-enriched FC at rest. In the first step, the fMRI signal in the gray matter was weighted using the molecular PET templates as a spatial regressors to estimate the dominant blood-oxygen-level-dependent (BOLD) fluctuations of the 5-HTT- and 5-HT1AR-enriched functional systems at the subject level. The cerebellum was excluded at this stage as it was used as a reference region in the kinetic model for both systems (Beliveau et al., 2017). In the second step, the subject-specific time series estimated in the first step were used as temporal regressors to estimate the subject-specific spatial maps of 5-HTT- and 5-HT1AR-enriched FC.

**Figure 2.**
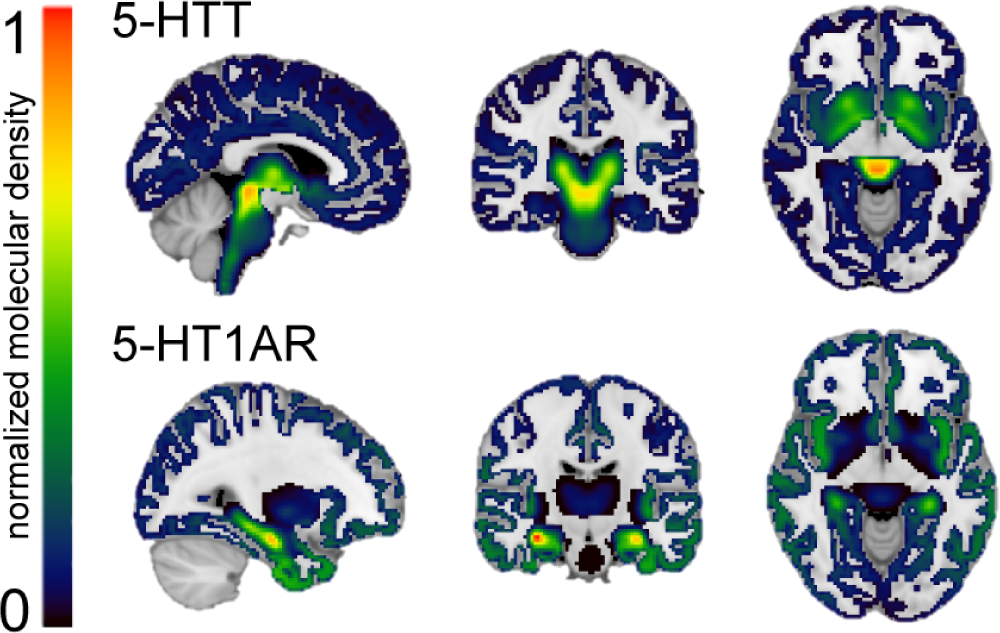
Normalized high-resolution in vivo Positron Emission Tomography (PET) maps of the distribution density of the serotonin transporter (5-HTT) and serotonin 1A receptor (5-HT1AR) 5-HTT and 5-HT1AR density distributions are normalized between 0 and 1 intensity (with 0 indicating a low density and 1 a high density) and overlaid onto a 1 mm Montreal Neurological Institution (MNI) template brain. The normalized distribution density of the 5-HTT is characterized by high density in the entorhinal and insular cortices, subcortical regions and the raphe nucleus (top). The normalized distribution density of the 5-HT1AR shows high density in the raphe nucleus, hippocampus, septum, amygdala and corticolimbic areas (bottom).

### Target-enriched functional response during the emotional face-matching task

For the tb-fMRI analysis, the dominant BOLD time series of the target-enriched functional systems were estimated at the subject level as described previously. In the second step, the resulting subject-specific time series were used to estimate the 5-HTT- and 5-HT1AR-enriched faces>shapes functional response using a generalized psychological-physiological interaction design with five regressors, namely the convolved faces>shapes regressor, combining both *faces* and *shapes* blocks (task effect), the BOLD time series of the 5-HTT- and 5-HT1AR-enriched functional systems (task-independent target-enriched FC), and the regressors of the interaction between the convolved faces>shapes regressor and these BOLD time series (task-dependent target-enriched FC).

### Statistical analyses

All voxelwise analyses were conducted using permutation tests in Randomise (5000 permutations; threshold-free cluster enhancement (Smith & Nichols, 2009); significance inferred when family wise error corrected p<0.05).

For the rs-fMRI analysis, we evaluated the voxel-wise correlation of the target-enriched FC maps with the individual thalamic [^123^I]FP-CIT BR. Subsequently, we conducted exploratory analyses to assess differences in the target-enriched FC maps between treatment groups using two-sample t-tests, applying a Bonferroni correction for multiple comparisons (three groups yielding four comparisons; significance inferred when p<0.0125).

For the tb-fMRI analysis, we investigated the voxel-wise correlation of the task activation map and target-enriched task-dependent FC maps with the individual thalamic [^123^I]FP-CIT BR. We also conducted exploratory analyses to assess group differences in task (de)activation and task-dependent FC. Here, we first identified clusters showing significant activity/connectivity in the placebo group using a one-sample t-test (to exclude drug-induced effects on the target-enriched FC). Using a small volume correction within these clusters, we subsequently assessed group differences using two-sample t-tests applying a Bonferroni correction for multiple comparisons. To assess the influence of our decision to use a small volume correction mask defined from the placebo group only, we conducted a sensitivity analysis where we first identified clusters showing significant activity/connectivity in all groups combined using a one-sample t-test, and then repeated the between-group comparisons as described above (Supplementary Results).

## Results

### Target-enriched FC at rest

The mean 5-HT1AR- and 5-HTT-enriched FC maps are visualized in Figure S1.

The correlation analyses revealed a significant negative relationship (*r* = −0.66) between the thalamic [^123^I]FP-CIT BR and 5-HTT-enriched FC, such that subjects with lower thalamic 5-HTT availability (high occupancy by citalopram) showed higher FC in some regions of the 5-HTT-enriched FC maps, including in the planum polare, central opercular cortex, temporal and occipital fusiform gyrus, temporal gyrus, and opercular cortex (Figure 3A). No significant relationship was found between thalamic [^123^I]FP-CIT BR and 5-HT1AR-enriched FC at rest.

**Figure 3.**
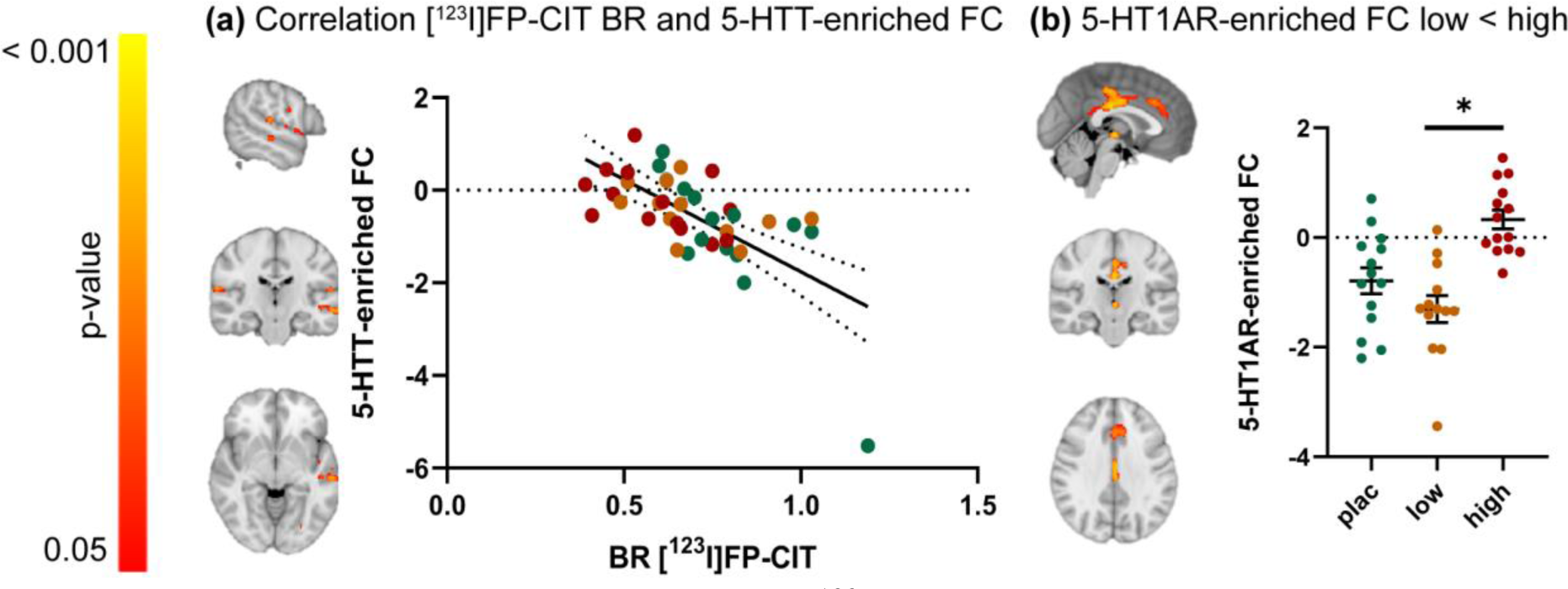
The relationship between thalamic [^123^I]FP-CIT ratio of specific to non-specific binding (BR) and serotonin transporter (5-HTT)- and serotonin 1A receptor (5-HT1AR)-enriched functional connectivity (FC) at rest. **(a)** Clusters showing a significant negative correlation between SPECT-derived thalamic [^123^I]FP-CIT BR and 5-HTT-enriched FC (left) and scatter dot plot illustrating this correlation (right). **(b)** Clusters showing significantly lower FC for the 5-HT1AR-enriched FC in the low dose group compared to the high dose group (left) (p < 0.05; FWE corrected) and scatter dot plot illustrating these results per treatment group (right). Error bars indicate mean±SEM. [^123^I]FP-CIT: [^123^I]N-ω-fluoropropyl-2β-carbomethoxy-3β-(4-iodophenyl)nortropane; low: low dose of citalopram (4 mg); high: high dose of citalopram (16 mg); SPECT: single-photon emission computerized tomography.

The exploratory analyses highlighted significant group differences in 5-HT1AR-enriched FC, such that FC of the low dose group in the posterior cingulate gyrus, anterior cingulate gyrus and thalamus was lower compared to the high dose group (Figure 3B). Further details on significant clusters can be found in Table S1. No significant pairwise group differences in 5-HTT-enriched FC were observed.

### Target-enriched functional response during the emotional face-matching task

The mean z-score maps of all task-based regressors are depicted in Figure S2.

We observed a significant negative association (*r* = −0.72) between the thalamic [^123^I]FP-CIT BR and task-dependent 5-HT1AR-enriched FC. Specifically, subjects with lower thalamic [^123^I]FP-CIT BR (i.e. high occupancy by citalopram) had higher FC in specific areas of the task-dependent 5-HT1AR-enriched maps, including the precentral gyrus, posterior cingulate and precuneus, opercular cortex, superior and middle frontal gyrus, frontal and temporal pole, temporal gyrus, thalamus, and left putamen (Figure 4A). We did not find a relationship between thalamic [^123^I]FP-CIT BR and task-dependent 5-HTT-enriched FC.

**Figure 4.**
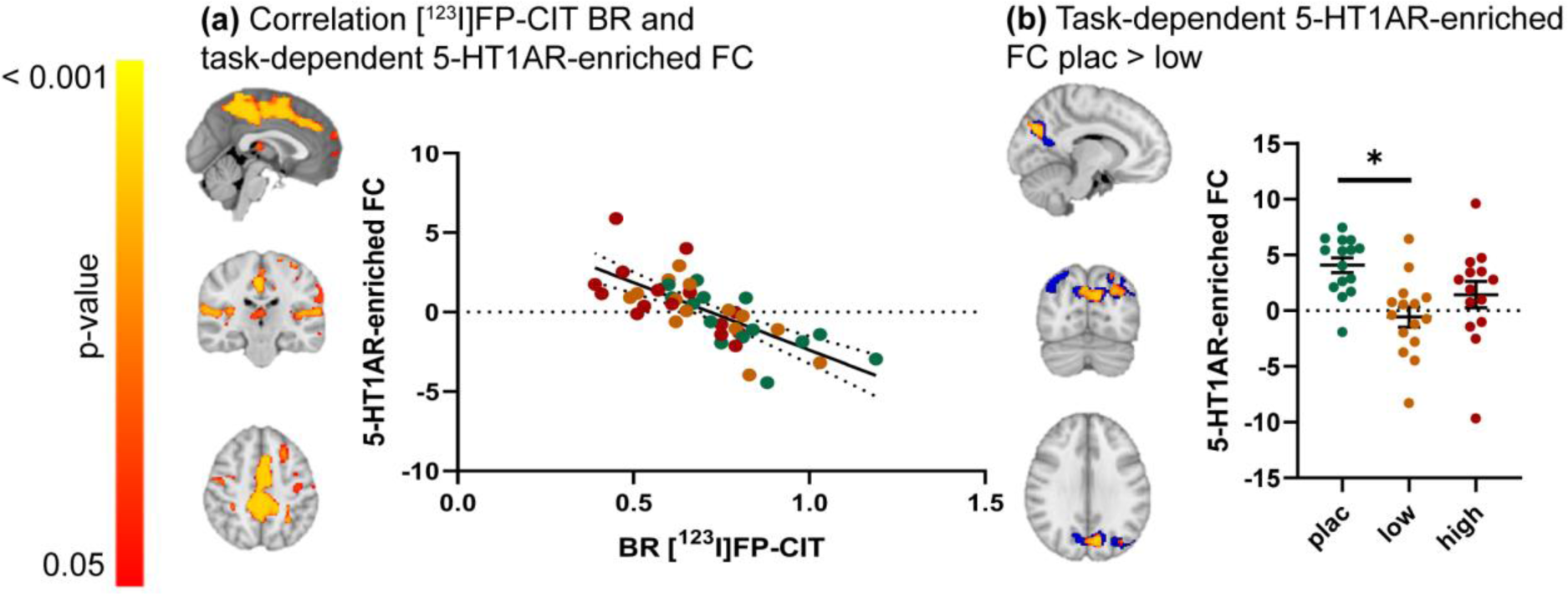
The relationship between thalamic [^123^]FP-CIT ratio of specific to non-specific binding (BR) and serotonin 1A receptor (5-HT1AR)-enriched task-dependent functional connectivity (FC) during an emotional face-matching task. **(a)** Clusters showing a significant negative correlation between SPECT-derived thalamic [^123^I]FP-CIT BR and task-dependent 5-HT1AR-enriched FC (left) and scatter dot plot illustrating this correlation (right). **(b)** Blue: clusters showing significant task-dependent 5-HT1AR-enriched FC, corresponding to the interaction between the convolved faces>shapes regressor and the dominant blood-oxygen level-dependent (BOLD) fluctuation related to the 5-HT1AR-enriched functional network, for the placebo group only (One sample t-test; p <0.05; FWE corrected). These areas were used as a small volume correction to constrain between-group comparisons. Red/yellow: clusters showing significant citalopram-induced dose-dependent differences in the 5-HT1AR-enriched task-dependent FC (p <0.05; FWE corrected) (left) and scatter plot depicting these group differences (right). Error bars indicate mean±SEM. [^123^I]FP-CIT:[^123^I]N-ω-fluoropropyl-2β-carbomethoxy-3β-(4iodophenyl)nortropane; plac: placebo; low: low dose of citalopram (4 mg); high: high dose of citalopram (16 mg); SPECT: single-photon emission computerized tomography.

The exploratory analysis showed a significant dose-dependent decrease in 5-HT1AR-enriched FC in the low dose group compared to the placebo group in the precuneus and lateral occipital cortex (Figure 4B). Cluster sizes and locations can be found in Table S2. No differences were found between the placebo and the high dose group, and between the low and the high dose groups. No group differences were observed in the task-dependent 5-HTT-enriched FC maps. Of note, the results of this exploratory analysis did not change significantly when using a small volume correction cluster derived from all groups combined (Figure S3 and Table S3).

No significant relationship was found between thalamic [^123^I]FP-CIT BR and task activation. The exploratory analysis did not show group differences in task activation.

## Discussion

In this study, we aimed to investigate how 5-HTT availability during an acute citalopram challenge is differentially associated with FC of 5-HTT- and 5-HT1AR-enriched networks at rest and during an emotional face-matching task. We found thalamic [^123^I]FP-CIT BR to be negatively associated with 5-HTT-enriched FC at rest, and with 5-HT1AR-enriched FC during an emotional face-matching task. Additionally, exploratory analyses revealed dose-dependent effects of citalopram on 5-HT1AR-enriched FC both at rest and during the task.

The (expected) negative relationship we observed between thalamic 5-HTT availability and 5-HTT-enriched FC showcases a direct link between citalopram-induced changes in neurotransmission and functional changes within a network defined by the canonical distribution of the drug’s main target in the healthy human brain. Therefore, our results align well with predictions from the simplest pharmacodynamic models that posit regional variability in 5-HTT density at the core of pharmacological effects on the brain. Interestingly, we observed that 5-HTT-enriched FC was less strong for individuals with lower 5-HTT availability in clusters encompassing the occipito-temporal and opercular cortex, which have previously been shown to exhibit alterations in connectivity and/or activity in patients with depression compared to healthy controls (Brown, Clark, Hassel, MacQueen, & Ramasubbu, 2017; Liu et al., 2014). The occipito-temporal cortex has been shown to respond to acute (but not chronic) treatment with escitalopram (Cheng et al., 2016). Contrastingly, we did not observe between-group differences in 5-HTT-enriched FC, which could reflect either a lack of power due to our modest sample size or unexplained across-group variability in 5-HTT availability that could hinder the identification of average group effects. For example, previous studies have highlighted gene polymorphisms in the 5-HTT-linked promoter region, which is known to affect functionality of the 5-HTT (Ruhé, Frokjaer, Haarman, Jacobs, & Booij, 2021).

In contrast to our hypothesis, we found no relationship between thalamic 5-HTT availability and 5-HT1AR-enriched FC at rest, suggesting that the 5-HT1AR-enriched functional response to citalopram is at least not linearly dependent on 5-HTT occupancy by citalopram. Interestingly, we found preliminary evidence of group differences in the PCC and anterior cingulate cortex (ACC), which are compatible with a nonlinear dose-response pattern where FC in the low dose group appeared lower in comparison with the placebo group but no differences between the high dose and placebo groups could be identified. However, we note that these findings should be interpreted with caution since they arise from exploratory pairwise tests in the absence of a significant overarching main effect of group. Moreover, interpreting this dose-response pattern on the 5-HT1AR-enriched functional network with our current understanding of the pharmacology of citalopram is admittedly challenging. One possible explanation could be that low and high doses of citalopram (indirectly) differently engage auto- and postsynaptic receptors. Here, a low dose could increase 5-HT to a smaller extent, and might predominantly affect auto- or postsynaptic 5-HT1AR; while the larger 5-HT increases induced by the high dose might engage both types of receptors, leading to compensatory effects summing to a null effect. Nevertheless, this hypothesis remains speculatory. It is worth noting that the regions where this effect occurred appear in line with previous literature and are highly relevant within the context of previous studies in depression. For instance, the PCC, as part of the DMN, has frequently been implicated in the functional response to SSRIs (Arnone et al., 2018; Klaassens et al., 2015; van de Ven et al., 2013; Van Wingen et al., 2014). The ACC is known to be densely innervated by serotonergic projections from the DRN, to have a relatively high density of 5-HT1AR and to show FC with both dorsal and medial raphe nucleus (Beliveau et al., 2015; Celada, Victoria Puig, & Artigas, 2013). Interestingly, a previous study suggested that ACC FC could serve as a predictor in escitalopram treatment outcome in patients (Tian et al., 2020), suggesting a crucial role for the ACC in the functional response to SSRIs. Whether this is mediated by the 5-HT1AR system remains to be elucidated.

In line with our hypothesis, results from the emotional face-matching task showed a negative relationship between 5-HTT availability and task-dependent 5-HT1AR-enriched FC. 5-HT1AR-enriched FC varied linearly with variations in 5-HTT availability in large clusters of cortical and subcortical areas, which show considerable overlap with the corticolimbic system. Interestingly, while previous studies have shown that limbic system activity during emotional processing is dependent on the 5-HT1AR (Fisher et al., 2006; Selvaraj et al., 2018), our findings suggest that 5-HT1AR-enriched FC in this network is dependent on 5-HTT availability. This emphasizes the important role for the 5-HT1AR in regulating fluctuating levels of extracellular 5-HT in these regions, in which previous studies have shown increased activation and decreased FC in response to emotional stimuli in depressed patients, as well as a normalization to levels similar to HC following SSRIs treatment (Delaveau et al., 2011; Wang, Hermens, Hickie, & Lagopoulos, 2012). Outside of the limbic system, our exploratory group-level analyses revealed a more complex non-linear dose-response pattern. In the precuneus, an integral component of the DMN which has previously been described to show alterations in activation and FC during emotion processing and following citalopram administration (Dutta et al., 2019; Yang, Tsai, & Li, 2020), we observe positive 5-HT1AR-enriched FC in the placebo group, which seems reduced in the low dose group and, to a lesser extent, in the high dose group. Taken together, our findings support the involvement of 5-HT1AR in shaping how increased levels of 5-HT induced by SSRIs modulate neural circuits involved in emotion processing, adding to the current enthusiasm around 5-HT1AR targeting compounds as promising therapeutic approaches.

We also investigated main effects of the task and treatment effect on BOLD responses, since this data has not been published previously. The main task effect we observed is in line with previous studies using a similar paradigm, showing increased BOLD signal mostly in key limbic areas (Del-Ben et al., 2005) and decreased BOLD signal in DMN areas (Fox et al., 2005). In contrast to prior literature, no effect of citalopram was observed on BOLD responses during the task contrast, adding to the mixed results from previous studies where an effect of citalopram has not been found consistently. For example, both decreased and increased amygdala activity following acute citalopram in HC was reported during fearful vs. neutral faces, respectively (Anderson et al., 2011; Selvaraj et al., 2018). Another study reported that acute citalopram administration enhanced activation of fusiform gyri and thalamus and attenuated response in right lateral OFC and right amygdala for aversive vs neutral faces in HC (Del-Ben et al., 2005). However, comparing these studies with ours is challenging, given the differences in citalopram dose and route of administration as well as analysis approach. For instance, all these prior studies used intravenous administration of either 7.5mg or 10mg of citalopram and used a ROI approach. It is possible that we missed small effects of citalopram because of the relatively low power of our study or because the effects of the acute oral and acute intravenous dose used by the previous studies are not comparable.

Citalopram is considered a conventional antidepressant medication, but several other compounds binding to 5-HT receptors have gained a renewed interest for the treatment of depression, including 3,4-methylenedioxymethamphetamine (MDMA) (Gill et al., 2020) and the psychedelic lysergic acid diethylamide (LSD) (Rossi, Hallak, Bouso Saiz, & Dos Santos, 2022). While these compounds have different binding profiles compared to citalopram, target-enriched FC analyses have recently helped to unravel potential overlap in 5-HT receptors involved in the functional response to these compounds. MDMA predominantly binds to 5-HTT (in addition to other monoamine transporters), but is a releasing agent in addition to a reuptake blocker (Oeri, 2021). Using REACT, MDMA administration was found to significantly modulate networks informed by the distribution of both 5-HTT and 5-HT1AR (Dipasquale et al., 2019). LSD shows a different pattern of affinity, with high agonist activity at several 5-HT receptors, as well as at dopaminergic D1 and D2 receptors (De Gregorio, Comai, Posa, & Gobbi, 2016). While not investigating the 5-HTT itself, Lawn et al., (2022) showed that LSD significantly affected FC within serotonin 1A, 1B, 2A, and dopamine D1 and D2 receptor-related networks. Interestingly, despite being a direct agonist to the 5-HT1AR, the 5-HT1AR-enriched maps showed only a very localized FC increase, while widespread effects in the serotonin 1B- and dopamine D1 receptor-enriched maps were observed. Therefore, alongside these previous studies, our data supports the idea that REACT has potential in aiding the dissection of brain pharmacodynamics of compounds targeting the 5-HT system. However, as the above-mentioned studies were conducted in healthy volunteers, future work including patients with depression would be of interest as suggested differences in the baseline function of the 5-HT system (including the 5-HT1A autoreceptor) between patients and controls might lead to different drug-related functional outcomes (Boldrini, Underwood, Mann, & Arango, 2008).

This study has some limitations. First, the sample included only a relatively small sample of healthy young women, which limits statistical power and impedes detection of smaller effects and the possibility to extrapolate findings beyond women. Therefore, future larger and well-powered studies attempting to replicate our findings would be welcome. Second, [^123^I]FP-CIT is a non-selective radioligand that binds to both the dopamine transporter and 5-HTT. However, since the thalamus is mostly devoid of dopamine transporters, thalamic binding provides the most reliable summary estimate of 5-HTT availability (Booij et al., 2007). Moreover, it should also be noted that the PET maps used to estimate the 5-HTT- and 5-HT1AR-enriched FC are average maps from PET images of healthy volunteers encompassing both men and women. It is unclear to what extent the distribution of the 5-HTT and 5-HT1AR is generalizable across genders. In addition, we did not have information about possible psychiatric disorders in first-degree relatives of our participants, although some 5-HT system alterations have tentatively been suggested for this group (Lochner, Chamberlain, Kidd, Fineberg, & Stein, 2016). Finally, as this study investigated only the effects of acute doses of citalopram, future studies should measure the functional response to citalopram at different time points to unravel its long-term effects on the 5-HTT and 5-HT1AR functional circuits.

## Conclusion

Building on the power of multimodal data acquired within the same participants, our study provides empirical evidence linking 5-HTT availability and FC of 5-HTT-enriched (at rest) and 5-HT1AR-enriched (during an emotional face-matching task) functional networks. Moreover, our study highlights the potential contribution of 5-HT1AR, in addition to 5-HTT, to dose-dependent changes in the functional response after acute citalopram administration. We provide tentative evidence for a complex dose-response pattern in healthy women, which differs between resting-state and an emotional face-matching task, and calls for further studies examining how dose, context and their interaction might moderate the effects of 5-HT-altering drugs on the brain. Our data supports the added value of REACT when studying the effects of citalopram and drugs alike on the brain. Given that the therapeutic effects of SSRIs are typically evaluated under chronic treatment, longitudinal studies examining 5-HTT- and 5-HT1AR-enriched FC changes over the course of treatment in patients, as well as its relationship with clinical response, would be interesting avenues to pursue in future research.

