## Supplementary Materials for "Modulation of functional networks related to the serotonin neurotransmitter system by citalopram: evidence from a multimodal neuroimaging study"

### SUPPLEMENTARY METHODS

#### MRI data preprocessing

Results included in this manuscript come from preprocessing performed using *fMRIPrep* 20.0.6 [1,2]; RRID:SCR_016216), which is based on *Nipype* 1.4.2 [3,4]; RRID:SCR_002502).

*Anatomical data preprocessing*

The T1-weighted (T1w) image was corrected for intensity non-uniformity (INU) with N4BiasFieldCorrection [5], distributed with ANTs 2.2.0 [6]; RRID:SCR_004757), and used as T1w-reference throughout the workflow. The T1w-reference was then skull-stripped with a *Nipype* implementation of the antsBrainExtraction.sh workflow (from ANTs), using OASIS30ANTs as target template. Brain tissue segmentation of cerebrospinal fluid (CSF), white-matter (WM) and gray-matter (GM) was performed on the brain-extracted T1w using fast (FSL 5.0.9, RRID:SCR_002823, [7]). Volume-based spatial normalization to two standard spaces (MNI152NLin2009cAsym, MNI152NLin6Asym) was performed through nonlinear registration with antsRegistration (ANTs 2.2.0), using brain-extracted versions of both T1w reference and the T1w template. The following templates were selected for spatial normalization: *ICBM 152 Nonlinear Asymmetrical template version 2009c* [8], RRID:SCR_008796; TemplateFlow ID: MNI152NLin2009cAsym], *FSL's MNI ICBM 152 non-linear 6th Generation Asymmetric Average Brain Stereotaxic Registration Model* [9], RRID:SCR_002823; TemplateFlow ID: MNI152NLin6Asym].

*Functional data preprocessing*

For each of the 1 BOLD runs found per subject (across all tasks and sessions), the following preprocessing was performed. First, a reference volume and its skull-stripped version were generated using a custom methodology of *fMRIPrep*. A deformation field to correct for susceptibility distortions was estimated based on *fMRIPrep*'s *fieldmap-less* approach. The deformation field is that resulting from co-registering the BOLD reference to the same-subject T1w-reference with its intensity inverted [10,11]. Registration is performed with antsRegistration (ANTs 2.2.0), and the process regularized by constraining deformation to be nonzero only along the phase-encoding direction, and modulated with an average fieldmap template [12]. Based on the estimated susceptibility distortion, a corrected EPI (echo-planar imaging) reference was calculated for a more accurate co-registration with the anatomical reference. The BOLD reference was then co-registered to the T1w reference using flirt (FSL 5.0.9, [13]) with the boundary-based registration [14] cost-function. Co-registration was configured with nine degrees of freedom to account for distortions remaining in the BOLD reference. Head-motion parameters with respect to the BOLD reference (transformation matrices, and six corresponding rotation and translation parameters) are estimated before any spatiotemporal filtering using mcflirt (FSL 5.0.9, [15]).The BOLD time-series (including slice-timing correction when applied) were resampled onto their original, native space by applying a single, composite transform to correct for head-motion and susceptibility distortions. These resampled BOLD time-series will be referred to as *preprocessed BOLD in original space*, or just *preprocessed BOLD*. The BOLD time-series were resampled into standard space, generating a *preprocessed BOLD run in MNI152NLin2009cAsym space*. First, a reference volume and its skull-stripped version were generated using a custom methodology of *fMRIPrep*. Automatic removal of motion artifacts using independent component analysis (ICA-AROMA, [16]) was performed on the *preprocessed BOLD on MNI space* time-series after removal of non-steady state volumes and spatial smoothing with an isotropic, Gaussian kernel of 6mm FWHM (full-width half-maximum). Corresponding “non-aggresively” denoised runs were produced after such smoothing. Additionally, the “aggressive” noise-regressors were collected and placed in the corresponding confounds file. Several confounding time-series were calculated based on the *preprocessed BOLD*: framewise displacement (FD), DVARS and three region-wise global signals. FD and DVARS are calculated for each functional run, both using their implementations in *Nipype* (following the definitions by [17]). The three global signals are extracted within the CSF, the WM, and the whole-brain masks. Additionally, a set of physiological regressors were extracted to allow for component-based noise correction (*CompCor*, [18]). Principal components are estimated after high-pass filtering the *preprocessed BOLD* time-series (using a discrete cosine filter with 128s cut-off) for the two *CompCor* variants: temporal (tCompCor) and anatomical (aCompCor). tCompCor components are then calculated from the top 5% variable voxels within a mask covering the subcortical regions. This subcortical mask is obtained by heavily eroding the brain mask, which ensures it does not include cortical GM regions. For aCompCor, components are calculated within the intersection of the aforementioned mask and the union of CSF and WM masks calculated in T1w space, after their projection to the native space of each functional run (using the inverse BOLD-to-T1w transformation). Components are also calculated separately within the WM and CSF masks. For each CompCor decomposition, the *k* components with the largest singular values are retained, such that the retained components’ time series are sufficient to explain 50 percent of variance across the nuisance mask (CSF, WM, combined, or temporal). The remaining components are dropped from consideration. The head-motion estimates calculated in the correction step were also placed within the corresponding confounds file. The confound time series derived from head motion estimates and global signals were expanded with the inclusion of temporal derivatives and quadratic terms for each [19]. Frames that exceeded a threshold of 0.5 mm FD or 1.5 standardised DVARS were annotated as motion outliers. All resamplings can be performed with *a single interpolation step* by composing all the pertinent transformations (i.e. head-motion transform matrices, susceptibility distortion correction when available, and co-registrations to anatomical and output spaces). Gridded (volumetric) resamplings were performed using antsApplyTransforms (ANTs), configured with Lanczos interpolation to minimize the smoothing effects of other kernels [20]. Non-gridded (surface) resamplings were performed using mri_vol2surf (FreeSurfer).

#### SUPPLEMENTARY RESULTS

*Mean maps of the target-enriched FC at rest*

The mean 5-HT1AR- and 5-HTT-enriched FC maps are visualized in Figure S1. The 5-HTT-enriched functional circuit involved the anterior cingulate, medial prefrontal cortex, juxtapositional lobule cortex, insular cortex, amygdala, thalamus, and the striatum (Figure S1A). The 5-HT1AR-enriched functional network comprised the temporal gyrus, hippocampus, anterior cingulate and paracingulate gyri, posterior cingulate and precuneus, inferior frontal gyrus, brain stem, and the angular gyrus (Figure S1B).

*Mean maps of the five task regressors of the task-based fMRI analyses*

Increased activation for the contrast faces>shapes was observed in the lateral occipital cortex, amygdala, thalamus, hippocampus, inferior and middle frontal gyrus, paracingulate gyrus, middle frontal/precentral gyrus, lateral occipital gyrus, insular cortex and superior parietal lobule was observed. Widespread deactivation was observed in task-negative areas, including the medial prefrontal cortex, precuneus, anterior and posterior cingulate, middle and superior temporal gyrus, and occipital fusiform cortex, and posterior parahippocampal gyrus (see Figure S2A).

The mean task-independent 5-HTT-enriched FC maps were very similar to the mean 5-HTT-enriched FC maps at rest, with positive FC in the anterior cingulate, medial prefrontal cortex, juxtapositional lobule cortex, insular cortex, amygdala, thalamus, and the striatum (Figure S2B) and negative 5-HTT-enriched FC in e.g. the paracingulate cortex, posterior cingulate cortex, occipital cortex, and parahippocampal gyrus. For the 5-HT1AR-enriched task-independent FC maps, we observed a positive FC with the bilateral hippocampus, fusiform cortex, parahippocampal gyrus, temporal gyrus, lateral occipital cortex, (para)cingulate gyrus and frontal pole and negative FC in e.g. the cingulate gyrus, precuneus, limbic system and occipital cortex independent of treatment group (Figure S2C).

The mean task-dependent 5-HTT-enriched FC maps show positive FC in the frontal gyrus, paracingulate gyrus, lateral occipital cortex, inferior temporal gyrus and brain stem, and negative FC in many cortical and subcortical regions (Figure S2D). The mean task-dependent 5-HT1AR-enriched FC maps show positive FC in e.g. posterior areas such as the posterior cingulate, precuneus, and the lateral occipital cortex, temporal areas such as the temporal fusiform cortex, parahippocampal gyrus, and hippocampus, frontal areas such as the paracingulate and cingulate gyri, and superior frontal gyrus, and the amygdala and thalamus during faces<shapes (Figure S2E).

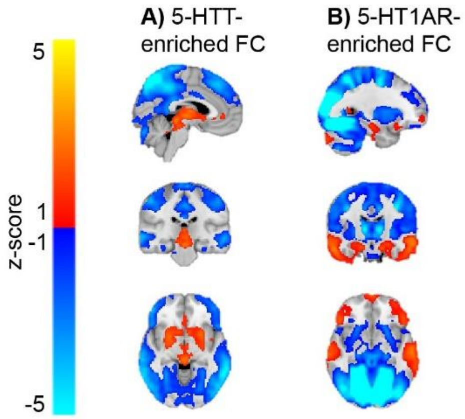


**Figure S1.** Mean maps of the serotonin transporter (5-HTT)- and serotonin 1A receptor (5-HT1AR)-enriched functional connectivity (FC) at rest

*The mean **A)** 5-HTT- and **B)** 5-HT1AR-enriched FC at rest was calculated by averaging spatial z-score maps for all subjects. Positive FC is depicted in red/yellow, whereas negative FC is depicted in blue ($p < 0.05$; FWE corrected).*

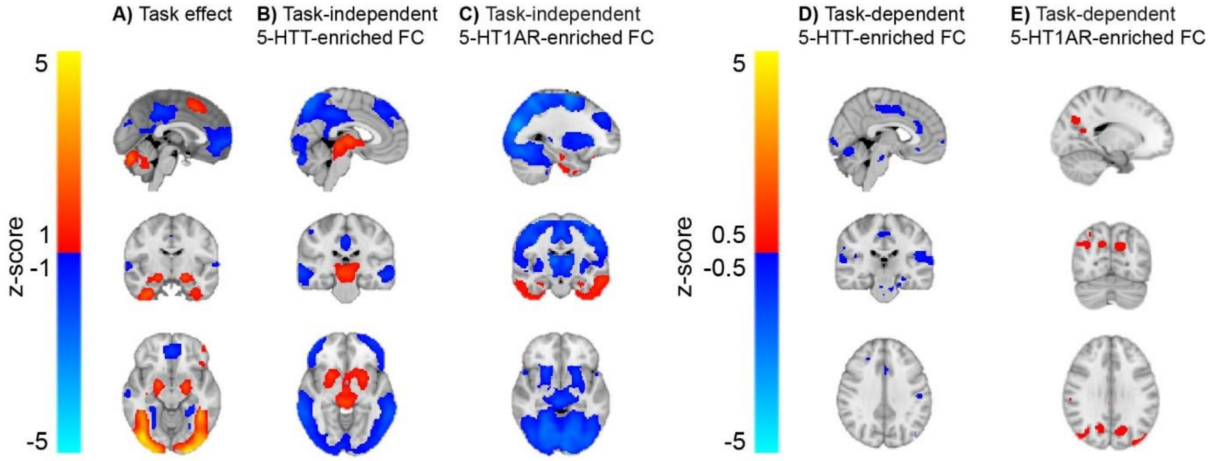


**Figure S2.** Mean z-score maps of the functional brain response during an emotional face matching task

***A)** The mean (de-)activation pattern associated with the faces>shapes task regressor. Activation is depicted in red/yellow, deactivation is depicted in blue. The mean task-independent **B)** 5-HTT- and **C)** 5-HT1AR-enriched functional connectivity (FC). The mean task-dependent target-enriched FC is depicted, corresponding to the interaction between the convolved faces>shapes regressor and the dominant blood-oxygen level-dependent (BOLD) fluctuation within the **D)** 5-HTT- and **E)** 5-HT1AR-enriched functional system. Positive FC during faces>shapes is depicted in red/yellow, negative FC is depicted in blue. Mean maps were constructed by calculating the mean z-score spatial maps across all subjects.*

**Table S1.** Clusters with significant group differences in 5-HTT- or 5-HT1A-derived functional connectivity for resting-state fMRI

*MNI: Montreal Neurological Institute; fMRI: functional MRI; rs-fMRI: resting-state fMRI; tb-fMRI: task-based fRMI; 5-HT: serotonin transporter; 5-HT1AR: serotonin 1A receptor; [$^{123}$I]FP-CIT: ; BP: binding potential; plac: placebo; low: low dose of citalopram (4 mg); high: high dose of citalopram (16 mg)*

| | | | | MNI coordinates | | | | |
|---|---|---|---|---|---|---|---|---|
| Modality | Result | Cluster | Size (voxels) | X | Y | Z | $t_{peak}$ | $p_{peak}$ |
| rs-fMRI | *5-HTT Correlation SPECT* | 1 | 164 | -55.2 | -1.81 | 1.57 | 5.63 | 0.028 |
| | | 2 | 131 | -22.8 | -67.3 | -14.3 | 6.57 | 0.016 |
| | | 3 | 122 | -56.9 | -24.4 | -6.29 | 6.57 | 0.017 |
| | | 4 | 52 | 62.2 | -27.8 | 15.9 | 6.73 | 0.02 |
| | | 5 | 46 | -56.9 | -24.6 | 13.4 | 6.44 | 0.023 |
| | | 6 | 34 | -41.1 | -27.8 | 15.9 | 5.6 | 0.033 |
| | | 7 | 26 | -59.2 | -3.46 | 23.8 | 5.41 | 0.04 |
| | | 8 | 23 | -61.6 | -11.7 | -9.13 | 4.55 | 0.046 |
| | | 9 | 20 | 35.9 | -29.6 | 18.2 | 6.39 | 0.031 |
| | *5-HT1AR low < high* | 1 | 129 | -0.635 | -20.7 | 33.9 | 6.01 | 0.007 |
| | | 2 | 12 | -1 | -18 | -3 | 7.36 | 0.008 |

**Table S2.** Clusters with significant group differences in 5-HTT- or 5-HT1A-derived functional connectivity for task-based fMRI during emotion recognition

*MNI: Montreal Neurological Institute; fMRI: functional MRI; rs-fMRI: resting-state fMRI; tb-fMRI: task-based fRMI; 5-HT: serotonin transporter; 5-HT1AR: serotonin 1A receptor; [123I]FP-CIT: ; BP: binding potential; plac: placebo; low: low dose of citalopram (4 mg); high: high dose of citalopram (16 mg)*

| | | | | MNI coordinates | | | | |
|---|---|---|---|---|---|---|---|---|
| Modality | Result | Cluster | Size (voxels) | X | Y | Z | $t_{peak}$ | $p_{peak}$ |
| tb-fMRI | *5-HT1AR Correlation SPECT* | 1 | 6915 | -2.39 | -26.2 | 53.2 | 6.87 | < 0.001 |
| | | 2 | 1785 | 51.4 | -13.1 | 18.5 | 4.75 | 0.005 |
| | | 3 | 798 | -55.3 | -25.7 | 16.4 | 5.05 | 0.006 |
| | | 4 | 252 | -23.5 | 24.2 | 44.3 | 4.6 | 0.015 |
| | | 5 | 249 | -39.9 | -7.89 | 52.2 | 3.37 | 0.036 |
| | | 6 | 117 | -57.6 | 2.73 | 6.05 | 4.28 | 0.027 |
| | | 7 | 109 | 3.43 | -21.4 | 9.56 | 4.35 | 0.028 |
| | | 8 | 80 | -1.63 | 62.6 | 23.7 | 4.15 | 0.031 |
| | | 9 | 71 | -42.9 | 13.1 | -33.6 | 5.2 | 0.019 |
| | | 10 | 39 | -47 | 24.4 | -8.25 | 4.07 | 0.041 |
| | | 11 | 36 | -23.8 | 45.4 | 32.3 | 3.37 | 0.045 |
| | | 12 | 26 | -0.692 | 62.2 | 3.15 | 3.76 | 0.045 |
| | | 13 | 24 | -63 | -18.7 | -14 | 3.12 | 0.045 |
| | | 14 | 7 | -24.3 | 0.857 | 12.3 | 3.65 | 0.047 |
| | | 15 | 4 | -22.5 | -5 | -37.5 | 4.47 | 0.049 |
| | *5-HT1AR plac > low* | 1 | 237 | -4.82 | -75.5 | 32.7 | 4.2 | 0.003 |
| | | 2 | 67 | -30.3 | -71.8 | 36 | 3.47 | 0.011 |
| | | 3 | 1.8 | -61 | 16.7 | 39.2 | 3.79 | 0.001 |

*Small volume correction based on significant activation in all treatment groups for the task-based fMRI analyses*

We additionally conducted a sensitivity analysis to assess the influence of our decision to estimate the significant 5-HTT- and 5-HT1AR-enriched FC associated with the regressors of our gPPI model. The mask used to constrain the permutation tests in Randomise (t-tests, corrected for multiple comparisons using a Bonferroni correction) is depicted in Figure S2A.

For the task regressor, we did not observe any group differences, in line with our previously described results. However, for the task-dependent 5-HT1AR-enriched FC, a group difference was detected, where the FC with the precuneus was higher for the placebo group compared to the (Figure S2 and Table S2).

All observed (absence of) group differences, as well as the location of the clusters showing significant group differences in the task-dependent 5-HT1AR-enriched FC, is highly similar to the results provided in the full manuscript. We therefore do not expect that our decision to constrain the permutation-based t-tests to the voxels showing significant FC in the placebo group only, has influenced the results significantly.

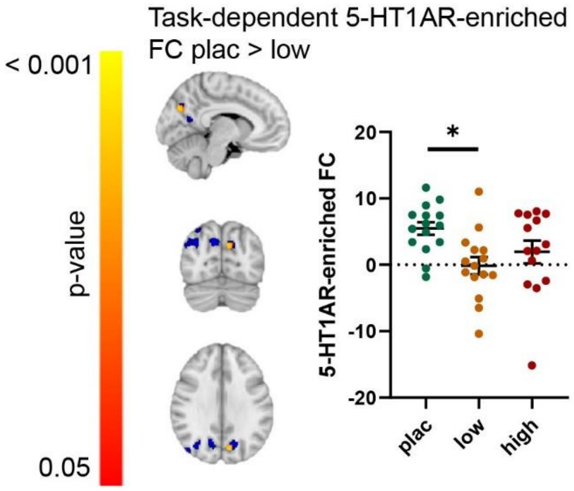


**Figure S3.** Citalopram-induced differences in task-dependent serotonin 1A receptor (5-HT1AR)-enriched functional connectivity (FC)

*Blue: clusters showing significant 5-HT1AR-enriched task-dependent FC, corresponding to the interaction between the convolved faces>shapes regressor and the dominant blood-oxygen level-dependent (BOLD) fluctuation related to the 5-HT1AR-enriched functional network, for all groups combined (One sample t-test; p < 0.05; FWE corrected). These areas were used as a small volume correction to constrain between-group comparisons. Red/yellow: clusters showing significant citalopram-induced dose-dependent differences in the 5-HT1AR-enriched FC maps. Error bars indicate mean±SEM.*

*plac: placebo; low: low dose of citalopram (4 mg); high: high dose of citalopram (16 mg)*

**Table S3.** Clusters with significant between-group task-dependent serotonin 1A receptor-enriched functional connectivity for task-based fMRI during an emotional face matching task

*MNI: Montreal Neurological Institute*

| | | MNI coordinates | | | |
|---|---|---|---|---|---|
| Cluster | Size (voxels) | X | Y | Z | $p_{peak}$ |
| 1 | 36 | -9.55 | -73.7 | 30.3 | 0.007 |
